# Global view on virus infection in non-human primates and implication for public health and wildlife conservation

**DOI:** 10.1101/2020.05.12.089961

**Authors:** Zhijin Liu

## Abstract

The pandemic outbreak and rapid worldwide spread of severe acute respiratory syndrome coronavirus 2 (SARS-CoV-2) is not only a threat for humans, but potentially also for many animals. Research has revealed that SARS-CoV-2 and other coronaviruses have been transmitted from animals to humans and *vice versa*, and across animal species, and hence, attracted public attention concerning host-virus interactions and transmission ways. Non-human primates (NHPs), as our evolutionary closest relatives, are susceptible to human viruses, and a number of pathogens are known to circulate between humans and NHPs. Here we generated global statistics of virus infection in NHPs (VI-NHPs). In total, 121 NHP species from 14 families have been reported to be infected by 139 DNA and RNA viruses from 23 virus families; 74.8 percent of viruses in NHPs have also been found in humans, indicative of the high potential for cross species transmission of these viruses. The top ten NHP species with high centrality in the NHP-virus network are two apes (*Pan troglodytes, Pongo pygmaeus*), seven Old World monkeys (*Macaca mulatta, M. fascicularis, Papio cynocephalus, Lophocebus albigena, Chlorocebus aethiops, Cercopithecus ascanius, C. nictitans*) and a lemur (*Propithecus diadema*). Besides apes, there is a high risk of virus circulation between humans and Old World monkeys, given the wide distribution of many Old World monkey species and their frequent contact with humans. We suggest epidemiological investigations in NHPs, specifically in Old World monkeys with close contact to humans, and other effective measures to prevent this potential circular transmission.

## Introduction

Coronavirus disease 2019 (COVID-19), caused by severe acute respiratory syndrome coronavirus 2 (SARS-CoV-2) rapidly spread worldwide, and recent studies suggest that pets and other animals could also be infected by SARS-CoV-2 through natural contact [1, 2]. Captive rhesus macaques (*M. mulatta*), inoculated with SARS-CoV-2 in pathological studies, exhibited a moderate infection as observed in the majority of human cases [3, 4]. Besides captive animals and pets, wild animals are also susceptible to the infection of coronaviruses transmitted from humans. For instance, in 2016, wild chimpanzees in Côte d’Ivoire were infected by the human coronavirus OC43 [5].

The close evolutionary relationship between humans and NHPs is thought to support pathogen transmission [6] and many viruses have been described that circulate between humans and NHPs. In captive and wild NHPs, various viruses including coronaviruses, enteroviruses, enteric adenoviruses, rotaviruses, and picobirnaviruses have been detected, which are also found in humans [7–9]. The most prominent cases of virus transmission from wild NHPs to human are simian foamy virus (SFV), yellow fever virus (YFV), Zika virus (ZIKV), and human immunodeficiency virus (HIV) [10–13]. Conversely, viruses such as poliovirus and measles have been reported in NHPs and likely derived from local human populations [14]. To block the potential circular transmission route of different viruses between human and NHPs, precautions and regulations are needed.

Here we performed a survey on documented virus infections in NHPs (VI-NHPs) based on published data. First, we generated a summary statistics of worldwide reported VI-NHPs. We then identified and predicted NHP species with a high risk of virus transmission from humans and predicted geographic locations where disease outbreaks are likely to occur.

## Materials and Methods

Global information of VI-NHPs was extracted from the Global Mammal Parasite Database (GMPD, http://www.mammalparasites.org/). We also used literature searches for publications describing VI-NHPs, which were not included in GMPD. Only the natural virus infections in captive and wild NHPs have been recorded, while the virus inoculations for pathological studies are not included.

We then built host-virus ecological networks in which nodes represent NHPs that are linked through shared viruses. Since centrality in primate-virus networks could assess the potential for the circulation of viruses among NHPs and humans, we estimated the centrality using four metrics: strength degree centrality, eigenvector centrality, betweenness centrality, and closeness centrality implemented in the R package “igraph” and UCINET 6.689 [15]. Since each metrics presents different and complementary aspects of centrality, we tested the correlations among all four centrality indices. To obtain a clearer picture of the effect of centrality of each NHP species on transmission ability, we obtained a composite centrality that integrates the different and complementary aspects of the four centrality metrics by performing a principal component analysis (PCA) on the centrality index correlations [13]. The phylogenetic generalized least squares (PGLS) methods were used to test the relationship between centrality and the number of viruses reported in each NHP species, and the number of viruses in each NHP species that are also found in human [13,16].

## Results

In total, 1,435 records of VI-NHPs were obtained from the GMPD. Forty-three additional publications describing VI-NHPs, not included in GMPD, were integrated into the overall database. Thus, our final dataset contained 1,478 records, describing infections caused by 139 different viruses in 121 NHP species (14 families, 49 genera) globally (Figures 1a and 1b). The viruses infecting NHPs covered DNA and RNA viruses from 23 families: Adenoviridae (20 viruses), Herpesviridae (17), Flaviviridae (15), Bunyaviridae (14), Retroviridae (13), Paramyxoviridae (10), Togaviridae (9), Picornaviridae (8), Polyomaviridae (4), Caliciviridae (4), Rhabdoviridae (4), Filoviridae (3), Orthomyxoviridae (3), Reoviridae (3), Poxviridae (2), Papillomaviridae (2), Coronaviridae (2), Picobirnaviridae (1), Parvoviridae (1), Astroviridae (1), Hepadnaviridae (1), Anelloviridae (1), and Arteriviridae (1) (Figure 1c). Among the 139 viruses reported in NHPs, 104 viruses (74.8%) were shared between human and NHPs, indicating the high zoonotic potential of these viruses.

**Figure 1.**
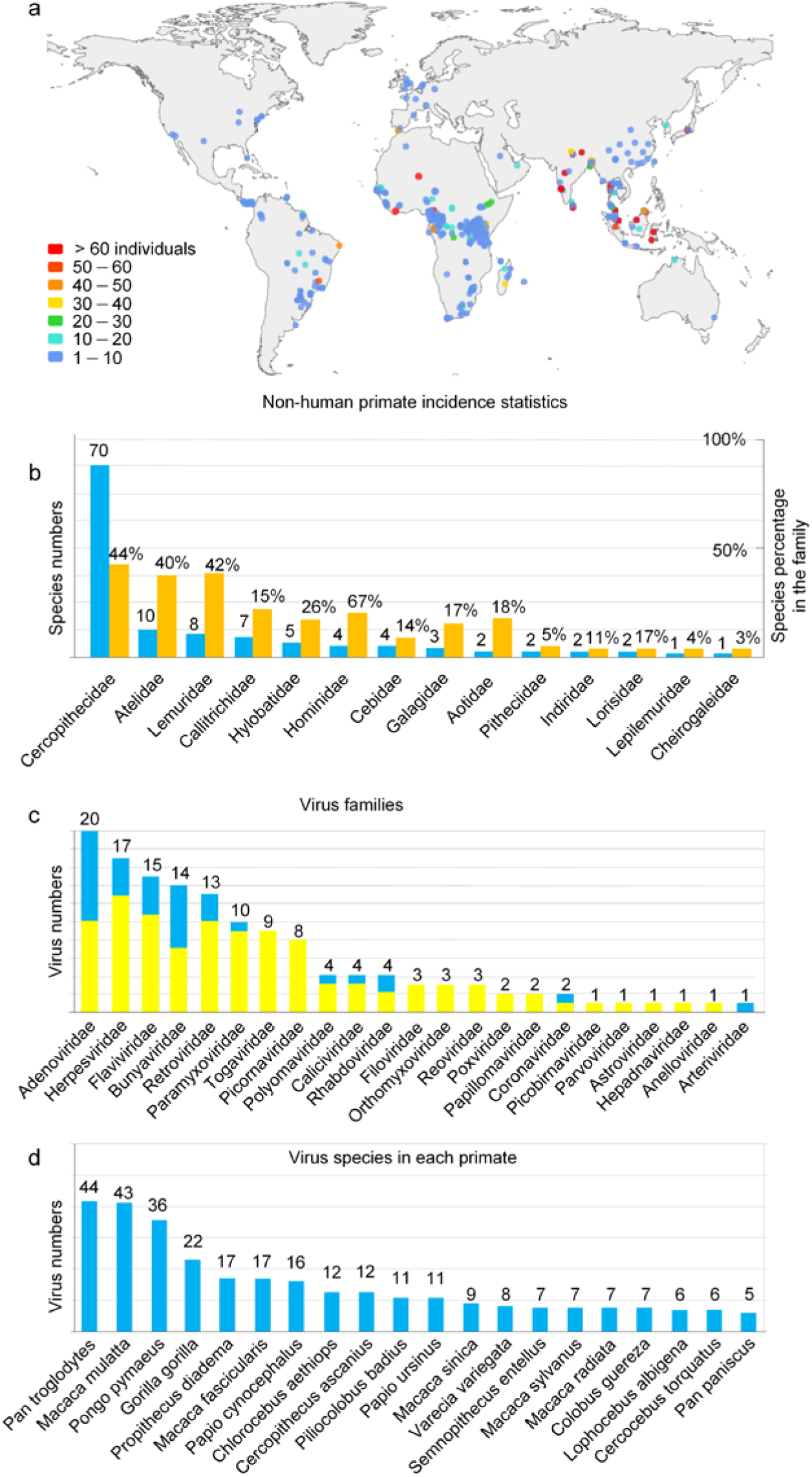
Incidence statistics for non-human primates. (a) Global patterns of known VI-NHPs, (b) NHP families infected by viruses, (c) virus species per virus family reported in NHPs, with yellow fraction referring to the number of viruses reported in humans, (d) number of viruses reported in NHPs (top 20).

The most documented VI-NHPs occurred in the chimpanzee (*Pan troglodytes*, 44 viruses; Figure 1d), but viruses were detected in all apes (chimpanzee, bonobo, gorillas and orangutans) and 94.4% of them have also been reported in humans. The second most documented VI-NHPs are found in the rhesus macaque (43 viruses; Figure 1d), of which 33 are shared with humans. Besides apes, Old World monkeys (Cercopithecidae) are with 70 (44.0%) infected species the most infected group among NHPs. Among other NHP families infected species range from one to ten (3.1-42.1%, Figure 1b).

We obtained a weighted unipartite NHP-virus network, in which each node represents a NHP species connected to other nodes by the edges weighted by the number of shared viruses (Figure 2a). All four centrality indices showed positive correlations (0.625 < *r* < 0.989, *P* < 0.0001 in all cases, n = 121 NHPs; Table S1), indicating that they detected similar NHP species as most central. A single factor found in PCA explained 85.2% of the variance of the indices, which was used as the composite index to assess the centrality of each node (Table S2). The top ten most central NHP species include two apes, seven Old World monkeys and a lemur, in descending order: *Pan troglodytes, Pongo pygmaeus, Papio cynocephalus, Macaca mulatta, Lophocebus albigena, Chlorocebus aethiops, Cercopithecus ascanius, Propithecus diadema, Macaca fascicularis*, and *Cercopithecus nictitans* (Figure 2a and 2b).

**Figure 2.**
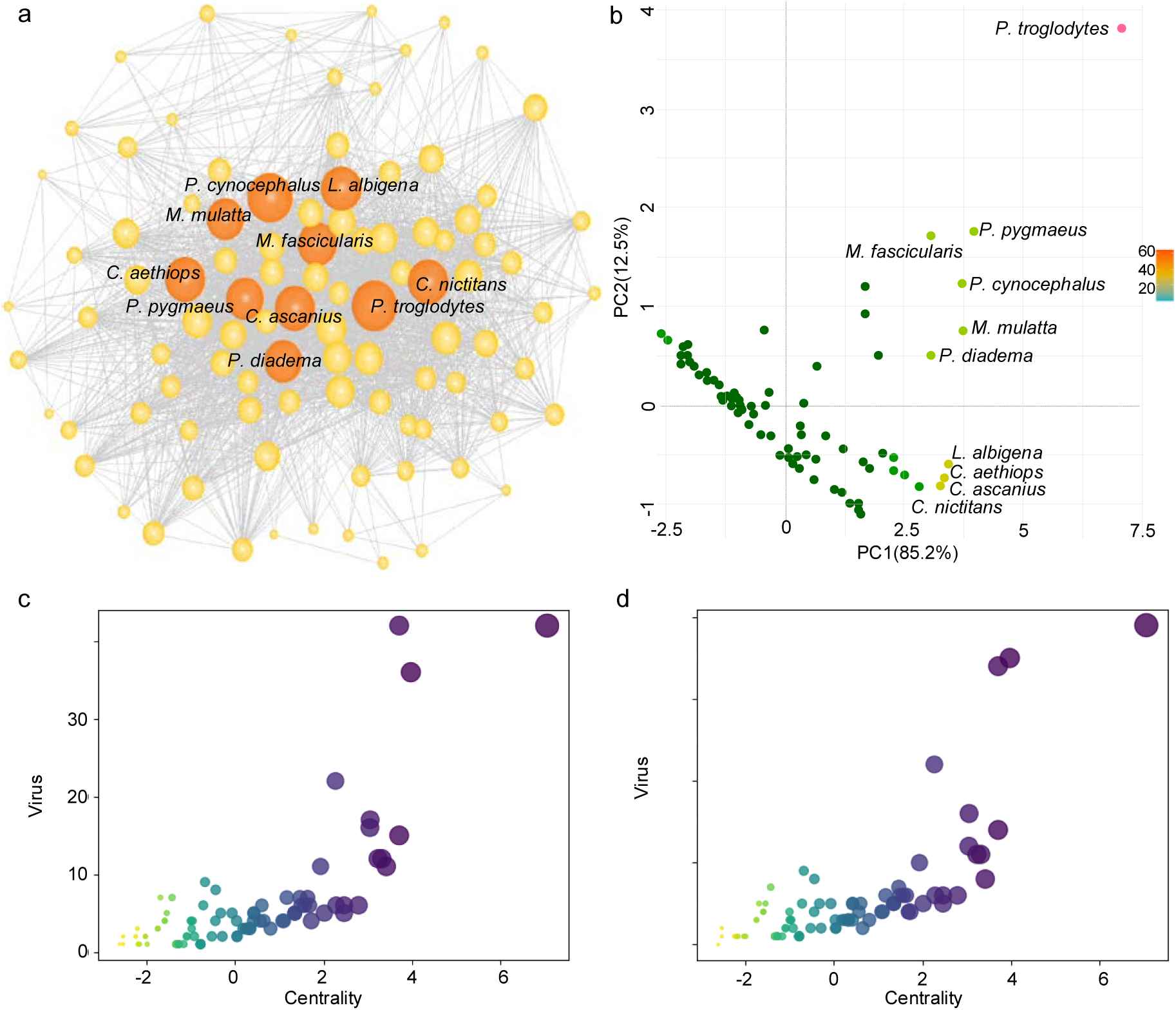
(a) Unipartite weighted network depicting the pattern of shared viruses by NHPs. Each node represents a NHP species. The links between nodes depict shared viruses (i.e., two nodes/species are linked whenever they share a virus). NHP species in the center of the network share more viruses than species on the periphery. The size of the nodes is proportional to the number of virus infections. (b) Composite index of centrality (PC1) of each NHP species in the network. (c) Relationship between centrality and the number of viruses in each NHP. (d) Relationship between centrality and the number of viruses that was also reported in humans in each NHP.

After controlling for phylogeny, virus number in each NHP species and the number of viruses shared with humans in each NHP species were significantly and positively associated to the centrality of each NHP species (strength degree centrality, eigenvector centrality, betweenness centrality, closeness centrality, and the composite centrality; Figure 2c and 2d, Table S3 and S4), respectively. This indicates that the centrality of a primate in the NHP-virus network could reflect the level of potential risk of virus transmission between NHPs and humans (and among NHPs).

## Discussion

Assuming that areas containing many NHP species and species evolutionary closely related to humans are more likely sources of zoonoses than areas containing fewer and more distantly related species, it was hypothesized that forests of central and western Africa represent areas where zoonotic outbreaks are likely to occur [17, 18]. Our study supports this hypothesis and suggests that African Old World monkeys (*P. cynocephalus, L. albigena, C. aethiops, C. ascanius*, and *C. nictitans*) exhibit a high potential for the circulation of viruses among NHPs and humans based on the centrality evaluation of our NHP-virus network. Interestingly, based on our statistics, the Asian rhesus macaque is the NHP species with the most reported virus infections. Rhesus macaques are, besides humans, the world’s most widely distributed primates, occupying a vast geographic distribution spanning from Afghanistan to the Chinese shore of the Pacific Ocean and south into Myanmar, Thailand, Laos, and Vietnam [19-22]. Furthermore, long-tailed macaques (*M. fascicularis*) are distributed over large parts of the Southeast Asian mainland and the Sundaland region between ca. 20°N and 10°S [23]. The network analyses shows the high centrality of these two macaque species in the NHP-virus network and ranked them among the top ten most central NHPs. Given the wide range of both macaque species, the centrality of macaques in the NHP-virus network and the frequent interface with humans (Figure 3a and 3b), there is a high risk of virus circulation between humans and macaques.

**Figure 3.**
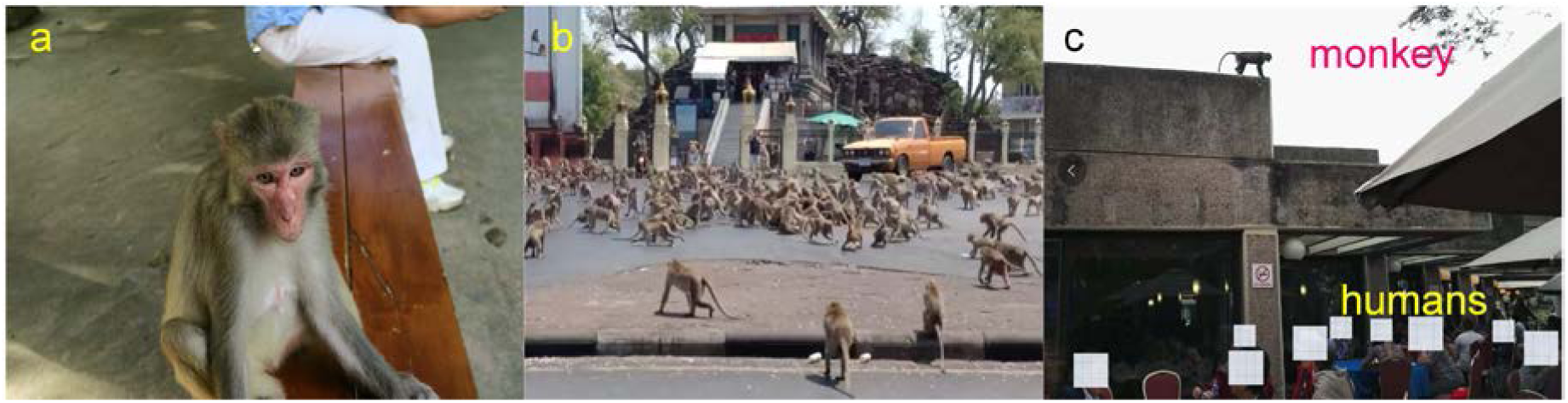
Close contact between NHPs and humans. (a) rhesus macaque (*M. mulatta*) in a city park (photographed by Bojun Liu), (b) long-tailed macaques (*M. fascicularis*) in a city (photo from website), and (c) blue monkey (*C. mitis*) watching people during the congress of the International Primatological Society 2018.

A major drawback of our study might be uneven and incomplete data as only few wild NHP populations have been thoroughly sampled. Since our statistics are based on documented VI-NHPs, records on virus infections are likely to be more complete and extensive for well-studied compared to less investigated NHP species. Centrality may also be affected by the number of studies on each NHP species, because more thoroughly sampled NHP species seem to have more confirmed virus infections [24]. However, there are also more interfaces between thoroughly studied NHPs and humans, which might lead to a higher probability of potential transmission. Thus, we did not control for the sampling effort in above analysis. For the sake of preciseness, we used the number of citations (=number of studies) as an estimate of sampling effort for each primate, including sampling effort in the computation of centrality estimates by upweighting the least sampled primates and down-weighting the most sampled primates [13]. Results showed the some trend with the analysis without controlling the sampling efforts, and for the sake of brevity we provide results in the supplementary metarials (Table S5-S9 and Figure S1). In the future, more efforts ought to be made for the collection, documentation and analysis of VI-NHP, especially for NHP species with higher potential of virus transmission. Since coronaviruses have been reported in macaques and other primates [7, 8], viral surveys should first target such species, not only to find known coronaviruses in such populations, but also to find new strains with high zoonotic potential.

Experts in animal health and conservation are starting to urge for the protection of great apes during human COVID-19 pandemics, since the transmission of the human virus to apes could result in severe outbreaks and local extinctions [25]. We suggest to expand such efforts to various Old World monkeys, as many of them, for instance, baboons or macaques, are widely distributed and often in close proximity to humans (Figure 3a, 3b and 3c). Moreover, bioinformatics analysis indicate that Old World monkeys, besides humans and apes, are more likely to be susceptible to SARS-CoV-2 than New World monkeys or strepsirrhines [26] and the rhesus macaques were most susceptible to SARS-CoV-2 infection compared to other Old World and New World monkeys [27]. Based on the centrality evaluation of our NPH-virus network, several Old World monkeys are considered to be at great risk of cross-species transmission due to the high centrality in the network. Since the distributions of Old World monkeys, apes, and humans often overlap, monitoring and regulations ought to be issued to block this potential circulative transmission route for the protection of NHPs. Especially macaques are widely used animal models with large captive populations almost all over the world [28]. Moreover, many wild Old World monkey species are in close contact with humans in national parks and even in urban districts (Figure 3a, 3b and 3c). Based on the above, route surveillances are necessary for captive and wild Old World monkey populations in frequent contact with humans.

## Acknowledgements

The project was supported by the Strategic Priority Research Program of the Chinese Academy of Sciences (XDA23080201, XDB31000000 and XDA19050202), the National Natural Science Foundation of China (31821001) and National Key R&D Program of China (2016YFC0503200). The authors thank Qi Wu, Zhenglong Wang, Paul Garber and Martin Burrows for data analyses and the written use of English.

## Author Contributions

Z.L., C.R. and M.L. conceived and designed the project. X.Q., L.Z., J.Z. and Z.Y. managed the project.

ORCIDs: C.R.: 0000-0003-0190-4266

