## Supplemental Data 1 for "Global view on virus infection in non-human primates and implication for public health and wildlife conservation"

**Supplementary Materials**

### Results

**Table S1.** Matrix of Product - Moment correlation among the Centrality Indices. All correlations have *p*-values < 0.0001.

| | Closeness Centrality | Betweenness Centrality | Eigenvector centrality |
|---|---|---|---|
| Betweenness Centrality | 0.6245 | | |
| Eigenvector centrality | 0.9026 | 0.6467 | |
| Strength centrality | 0.9266 | 0.6810 | 0.9889 |

**Table S2.** Result of the Principal Component Analysis on the correlation among the Centrality Indices.

| | Factor 1 (Composite Centrality) | Factor 2 |
|---|---|---|
| Percent of explained variance | 0.851 | 0.1183 |
| Closeness Centrality | 0.512 | 0.276 |
| Betweenness Centrality | 0.424 | -0.904 |
| Eigenvector centrality | 0.524 | 0.256 |
| Strength centrality | 0.532 | 0.203 |

**Table S3.** Result of phylogenetic GLS testing the effect of each centrality index on the number of viruses reported in each NHP.

| | Estimate ± 1 SE | t | *p*-value | $R^2$ | Lambda |
|---|---|---|---|---|---|
| Closeness Centrality | 0.4477±0.0643 | 6.9659 | < 0.0001 | 0.3311 | 0.192* |
| Betweenness Centrality | 0.1095±0.0061 | 17.9201 | < 0.0001 | 0.7693 | 0.000 |
| Eigenvector Centrality | 24.1574±2.1502 | 11.2350 | < 0.0001 | 0.5661 | 0.354*** |
| Strength Centrality | 0.1135±0.0100 | 11.3226 | < 0.0001 | 0.5699 | 0.351*** |
| Composite Centrality | 3.3226±0.2631 | 12.6284 | < 0.0001 | 0.6228 | 0.304*** |

Significance codes for phylogenetic signal (λ): * $p < 0.05$; *** $p < 0.001$.

**Table S4.** Result of phylogenetic GLS testing the effect of each centrality index on the number of viruses reported in each NHP shared by humans.

| | Estimate ± 1 SE | t | *p*-value | $R^2$ | Lambda |
|---|---|---|---|---|---|
| Closeness Centrality | 0.4099±0.0579 | 7.0824 | < 0.0001 | 0.3387 | 0.181 |
| Betweenness Centrality | 0.0982±0.0057 | 17.1101 | < 0.0001 | 0.7524 | 0.000 |
| Eigenvector Centrality | 22.0750±1.9301 | 11.4372 | < 0.0001 | 0.5749 | 0.351*** |
| Strength Centrality | 0.1034±0.0090 | 11.4533 | < 0.0001 | 0.5756 | 0.339*** |
| Composite Centrality | 3.0201±0.2374 | 12.7203 | < 0.0001 | 0.6262 | 0.294*** |

Significance codes for phylogenetic signal (λ): *** $p < 0.001$.

**Controlling for sampling bias in centrality computation.**

Host-virus data are sensitive to sampling effort. Consequently, the computation of individual centralities is largely influenced by the intensity of sampling of each primate species. We dealt with this issue by up-weighting the least sampled primate and down-weighting the most sampled primate per edge. For this, we corrected the weight of each edge by:

Sharing Viruses * 1/(Sampling Effort Primate 1* Sampling Effort Primate 2/Mean (Sampling Effort))

where Sampling Effort are the number of studies for each primate species.

All four centrality indices showed positive correlations ($0.2295 < r < 0.9723$, $P < 0.05$ in all cases, n = 121 NHPs; Table S5), indicating that they detected similar NHP species as most central. A single factor found in PCA explained 62.2% of the variance of the indices, which was used as the composite index to assess the centrality of each node (Table S2).

**Table S5.** Matrix of Product - Moment correlation among the Centrality Indices. All correlations have *p*-values < 0.05.

| | Closeness Centrality | Betweenness Centrality | Eigenvector centrality |
|---|---|---|---|
| Betweenness Centrality | 0.5372 | | |
| Eigenvector centrality | 0.2598 | 0.4115 | |
| Strength centrality | 0.2295 | 0.4724 | 0.9723 |

**Table S6.** Result of the Principal Component Analysis on the correlation among the Centrality Indices.

| | Factor 1 (Composite Centrality) | Factor 2 |
|---|---|---|
| Percent of explained variance | 0.622 | 0.264 |
| Closeness Centrality | 0.357 | 0.701 |
| Betweenness Centrality | 0.469 | 0.437 |
| Eigenvector centrality | 0.568 | -0.401 |
| Strength centrality | 0.575 | -0.396 |

**Table S7.** Result of phylogenetic GLS testing the effect of each centrality index on the number of viruses reported in each NHP.

| | Estimate ± 1 SE | t | *p*-value | $R^2$ | Lambda |
|---|---|---|---|---|---|
| Closeness Centrality | 3.9944±0.6121 | 6.5255 | < 0.0001 | 0.3022 | 0.000 |
| Betweenness Centrality | 0.0321±0.0054 | 5.8565 | < 0.0001 | 0.7693 | 0.000 |
| Eigenvector Centrality | 7.2118±2.8950 | 2.4911 | < 0.05 | 0.0514 | 0.000 |
| Strength Centrality | 0.1898±0.0731 | 2.5981 | < 0.05 | 0.0565 | 0.000 |
| Composite Centrality | 2.1307±0.4197 | 5.0773 | < 0.0001 | 0.2052 | 0.000 |

Significance codes for phylogenetic signal (λ): * $p < 0.05$.

**Table S8.** Result of phylogenetic GLS testing the effect of each centrality index on the number of viruses reported in each NHP shared by humans.

| | Estimate ± 1 SE | t | *p*-value | $R^2$ | Lambda |
|---|---|---|---|---|---|
| Closeness Centrality | 3.7336±0.5466 | 6.8301 | < 0.0001 | 0.3223 | 0.000 |
| Betweenness Centrality | 0.0305±0.0048 | 6.2737 | < 0.0001 | 0.2855 | 0.000 |
| Eigenvector Centrality | 7.1775±2.6056 | 2.7547 | < 0.001 | 0.5749 | 0.000 |
| Strength Centrality | 0.1890±0.0657 | 2.8769 | < 0.001 | 0.0642 | 0.000 |
| Composite Centrality | 2.0553±0.3733 | 5.5055 | < 0.0001 | 0.2339 | 0.000 |

Significance codes for phylogenetic signal (λ): * $p < 0.05$.

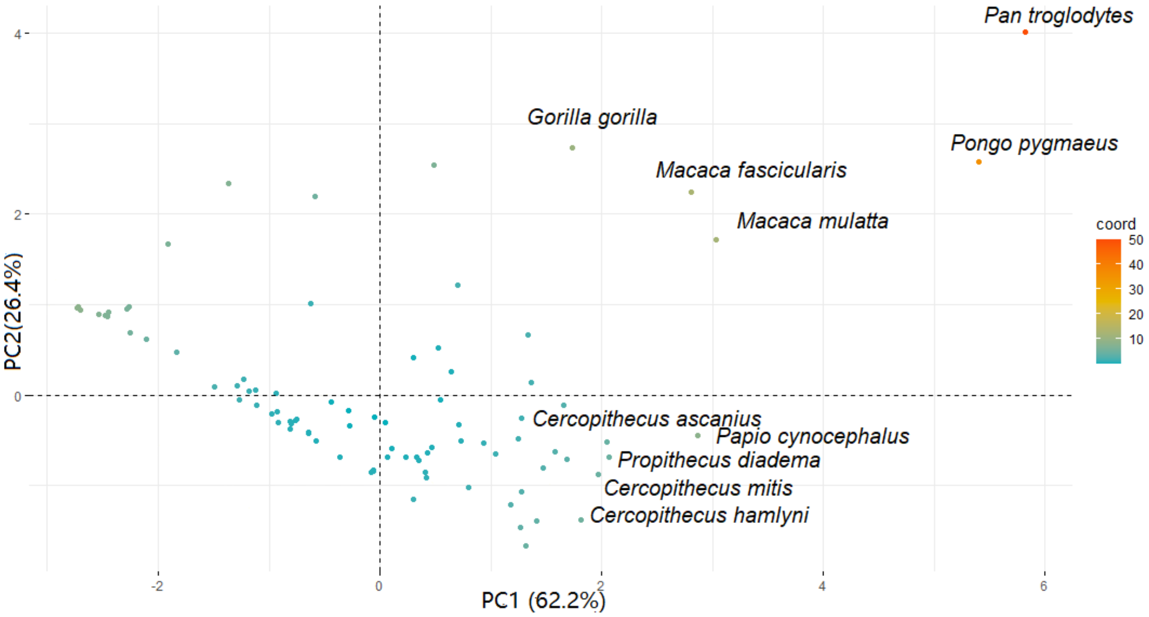


**Figure S1.** Composite index of centrality (PC1) of each NHP species in the network controlling the sampling efforts.

**Table S9.** The top ten of the most central primates generated by network analysis with and without controlling the sampling effort. In bold those species appearing in both lists.

| Rank | Not Controlling the sampling effort | Controlling the sampling effort |
|---|---|---|
| 1th | ***Pan troglodytes*** | ***Pan troglodytes*** |
| 2nd | ***Pongo pygmaeus*** | ***Pongo pygmaeus*** |
| 3rd | ***Papio cynocephalus*** | ***Macaca mulatta*** |
| 4th | ***Macaca mulatta*** | ***Papio cynocephalus*** |
| 5th | *Lophocebus albigena* | ***Macaca fascicularis*** |
| 6th | *Chlorocebus aethiops* | ***Propithecus diadema*** |
| 7th | ***Cercopithecus ascanius*** | ***Cercopithecus ascanius*** |
| 8th | ***Propithecus diadema*** | *Cercopithecus mitis* |
| 9th | ***Macaca fascicularis*** | *Cercopithecus hamlyni* |
| 10th | *Cercopithecus nictitans* | *Gorilla gorilla* |
